# Characterization of a Novel Type Homoserine Dehydrogenase Only with High Oxidation Activity from *Arthrobacter nicotinovorans*

**DOI:** 10.1101/2021.02.09.430557

**Authors:** Xinxin Liang, Huaxiang Deng, Yajun Bai, Tai-Ping Fan, Xiaohui Zheng, Yujie Cai

## Abstract

Homoserine dehydrogenase (HSD) is a key enzyme in the synthesis pathway of the aspartate family of amino acids. HSD can catalyze the reversible reaction of L-aspartate-β-semialdehyde (L-ASA) to L-homoserine (L-Hse). In direct contrast, growth characteristic studies of some bacterial such as *Arthrobacter nicotinovorans* showed that the bacterium could grow well in medium with L-homoserine as sole carbon, nitrogen and energy source, but the genes responsible for the degradation of L-Hse remain unknown. Based on the function and sequence analysis of HSD, one putative homoserine dehydrogenase from *A.nicotinovorans* was named AnHSD, which was different from those HSDs that from the aspartic acid metabolic pathway, might be responsible for the degradation of L-Hse. Surprisingly, the analysis showed that the purified AnHSD exhibited specific L-Hse oxidation activity without reducing activity. At pH 10.0 and 40 °C, The *K_m_* and *K_cat_* of AnHSD was 6.30 ± 1.03 mM and 462.71 s^-1^, respectively. AnHSD was partiality for NAD^+^ cofactor, as well as insensitive to feedback inhibition of downstream amino acids of aspartic acid family. The physiological role of AnHSD in *A.nicotinovorans* is discussed. These findings provide a novel insight for a better understanding of an alternative genetic pathway for L-Hse catabolism which was dominated by the novel HSD.

**Importance:** L-homoserine is an important building block for the synthesis of L-threonine, L-methionine, L-lysine which from aspartic acid family amino acids. However, some bacteria can make use of L-homoserine as a sole carbon and nitrogen source. Although the microbial degradation of L-homoserine has been studied several times, the genes involved and the molecular mechanisms remain unclear. In this study, we show that AnHSD responsible for the catabolism of L-homoserine in strain *Arthrobacter nicotinovorans*, as a special homoserine dehydrogenase with high diversity exists in *Arthrobacter*, *Microbacterium*, *Rhizobium*. We report for the first time that this novel homoserine dehydrogenase is now proposed to play a crucial role in that L-homoserine can use as a sole carbon and nitrogen source. This study is aimed at elucidating the enzymatic properties and function features of homoserine dehydrogenase from *Arthrobacter nicotinovorans*. These findings provide new insight into the catabolism of L-homoserine in bacteria.

## Introduction

Homoserine dehydrogenase (HSD; EC 1.1.1.3) exists in almost all plants and most microbes (1, 2), is a known NAD(P)H-dependent oxidoreductase that catalyzes the bidirectional reaction between L-aspartate-β-semialdehyde (L-ASA) and L-homoserine (L-Hse). HSDs from the aspartate metabolic pathways exhibit both oxidation and reduction activities, but its function tend to be more reduction for synthesizing L-Hse. L-Hse is a precursor for the synthesis of essential amino acids such as threonine, methionine, isoleucine in the L-aspartate family amino acids (AFAAs) (3, 4). According to the function and structure specificities, HSDs are classified into distinct families, namely, monofunctional HSDs and bifunctional AK-HSDs. For instance, HSDs from *Corynebacterium glutamicum* (CgHSD) and *Saccharomyces cerevisiae* (ScHSD) only exhibit monofunctional HSDs (5–7). Bifunctional HSDs are a fusion protein that monofunctional HSD fused aspartokinase (AK) at the N-terminal, like AK-HSDs from *Escherichia coli* (AK-HSDI and AK-HSDII) and *Arabidopsis thaliana* (AK-HSDI and AK-HSDII) (8, 9).

Several strains have been reported to partially or completely degrade L-Hse. *Rhizobium leguminosarum* reportedly uses L-Hse as its source of carbon and energy through the independent-aspartate metabolic pathway. Some putative genes (pRL80083, Rlv3841 and pRL80071) were found to be responsible for the catabolism of L-Hse in strain *R.leguminosarum* (10, 11). Mochizuki et al. reported the enantioselectively degradation of L-Hse from DL-homoserine by *Arthrobacter nicotinovorans* to obtain optically pure D-homoserine (12). However, the genes and enzymes responsible for the L-Hse biodegradation have seldom been reported in *A.Nicotinovorans*. We performed a genome-wide screen for essential genes in L-Hse metabolism from *A.Nicotinovorans* and found two putative HSD gene sequences. Based on the current reports, one protein AnHSD-109 (WP_055972109.1) was annotated as homoserine dehydrogenase and had a similar structure and function to CgHSD and ScHSD that were part of the L-aspartic acid pathway (13, 14). However, another protein AnHSD (WP_064723327.1) was not homologous to AnHSD-109. Therein, the similarity is measurably less than 40%, which means AnHSD may have special function and properties. To date, the researches on the structures, functions, and biochemical properties of this new type enzyme have not been reported in detail. In the present work, we reported a series of enzymatic properties of AnHSD with particular functions from *A. nicotinovorans*. The related research on the function of AnHSD may reveal a new metabolic pathway for microorganism to utilize L-Hse as carbon source.

## Results

### Identification and cloning of potential homoserine dehydrogenase

*A.Nicotinovorans* encodes a protein AnHSD of 348 amino acids and has a calculated molecular mass of approximately 35.97 kDa with a theoretical pI of 4.63 (http://www.expasy.ch/tools/protparam.html). A BLAST-P analysis (https://blast.ncbi.nlm.nih.gov/Blast.cgi) discloses many putative homoserine dehydrogenase from the genera *Arthrobacter*, and a few came from *Microbacterium, Streptomyces* and *Pseudomonas*. The top hits with characterized enzymes mostly involved *Staphylococcus aureus* (36.98% identity), *Thermus thermophilus HB8* (36.01% identity), *Sulfolobus tokodaii* (35.31% identity), *Hyperthermophilic archaeal* (31.75% identity). Identification of the protein family and protein domains was performed using an Interpro scan from EMBL-EBI (http://www.ebi.ac.uk/interpro/). This scan confirmed that AnHSD is a member of the homoserine dehydrogenase lacking ACT domain superfamily (IPR022697), containing an N-terminal NAD-binding homoserine dehydrogenase domain (IPR00001342) (amino acids [aa] 10 to 150), and an homoserine dehydrogenase domain (IPR005106) (aa 158 to 336).In addition, to identify and compare the conserved residues between AnHSD and other HSDs in more details, we retrieved 21 representative sequences from various species for sequence alignment analysis. Multiple sequence alignment revealed the conserved sequence motifs G-X-G-X-X-G/A/N was reported to be important for NAD(P)^+^ binding located at N-terminal (Fig.1A) (15), and the highly conserved sequences between 180 and 210 amino acids that were important for the catalytic activity of AnHSD (Fig.1B) (15–17).

**Figure 1.**
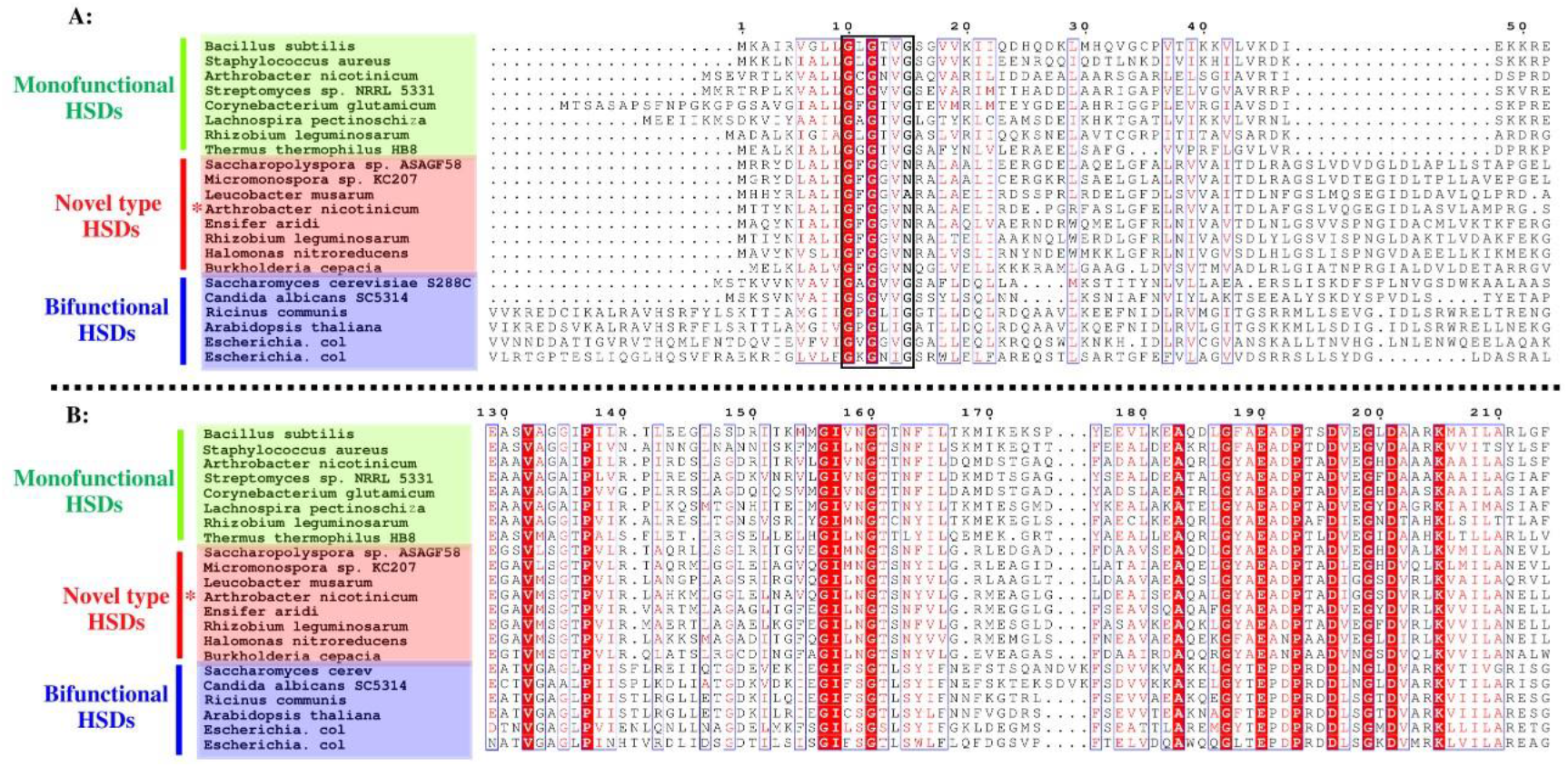
Sequence alignment. *Bacillus subtilis* (WP_014477878.1), *Staphylococcus aureus* (OHS88698.1), *Streptomyces sp. NRRL 5331* (CAC85208.1), *Corynebacterium glutamicum* (WP_003854900.1), *Lachnospira pectinoschiza* (WP_055174092.1), *Rhizobium leguminosarum* (which has two kinds of HSDs, WP_011654697.1 and WP_011651702.1), *Saccharopolyspora sp. ASAGF58* (WP_168589370.1), Micromonospora sp. KC207 (WP_132396959.1), Leucobacter musarum (WP_053351589.1), *Ensifer aridi* (WP_026617246.1), *Halomonas nitroreducens* (WP_053351589.1), *Burkholderia cepacian* (WP_027790144.1), *Saccharomyces cerevisiae S288C* (NP_012673.3), *Candida albicans SC5314* (XP_721450.1), *Ricinus communis* (EEF36869.1), *Arabidopsis thaliana* (NP_974576.1), *Escherichia coli str. K-12* (which has two kinds of HSDs, NP_414543.1 and NP_418375.1), *A. Nicotinovorans* (which has two kinds of HSDs, WP_064723327.1 and WP_055972109.1) were included in the alignment. Colored green for the monofunctional HSDs, red for a novel type HSDs, blue for bifunctional AK-HSDs. The red asterisk corresponds to the AnHSD (WP_064723327.1). (A). The black box for the NAD(P)-binding sites motif GXGXXG/A/N. (B). The highly conserved sequences between 180 and 210 amino acids for the catalytic activity sites of HSDs.

Some reported HSD sequences from other creatures were gathered to analyze their phylogenetic relationships. The phylogenetic tree was divided into three clusters representing bifunctional enzyme superfamily, monofunctional enzyme superfamily, and a novel type of HSDs (Fig.2). Enzymes in bifunctional HSDs superfamily exhibited the ability of AK and HSD activities, like AK-HSDs from *E.coli* and *A.thaliana*. CgHSD and BsHSD (which were from *Bacillus subtilis*) belonged to monofunctional HSD superfamiliy that showed HSD activity and did not have AK activity. Different from most reported bifunctional and monofunctional HSDs, almost no studies have been performed so far on the third type HSDs in terms of structure and function. Notably, Two homoserine dehydrogenase, AnHSD and AnHSD-109, from *A.nicotinophilus* were clustered into two branches of the phylogenetic tree, protein-sequence similarity was only 33.96% pairwise similarity. We analyzed that AnHSD and AnHSD-109 have evolved under different selective pressures, which caused AnHSD lost the reductive activity of L-ASA but with oxidation activity of L-Hse only.

**Figure 2.**
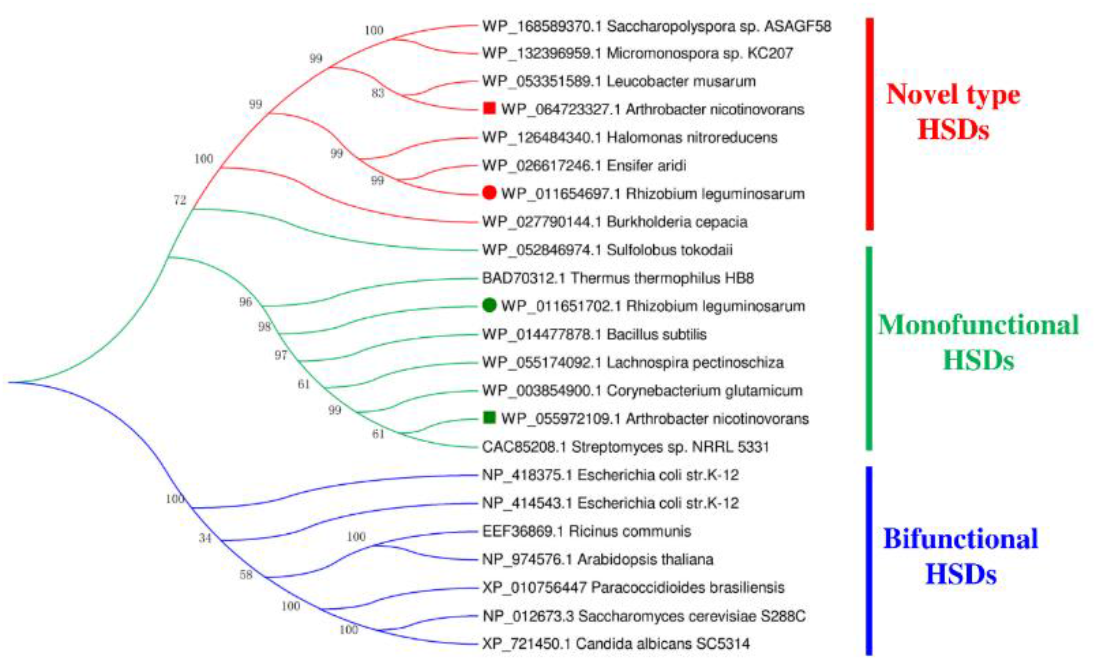
Phylogenetic tree for the HSDs from microorganism and plants. 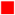 representing AnHSD of *A. nicotinovorans* (accession number WP_064723327.1); 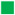 representing AnHSD-109 of *A. nicotinovorans* (accession number WP_055972109.1); 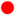 representing RlHSD-697 of *R. leguminosarum* (accession number WP_011654697.1); 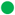 representing RlHSD-702 of *R. leguminosarum* (accession number WP_011651702.1).

### Purification of recombinant AnHSD and production determination

After transformation into *E.coli (DE3*) strains, the plasmids were validated by colonies PCR using primers specific for the expression vectors and flanking the cloning sites. The results indicated that AnHSD and pGro7 chaperone plasmid were successfully transformed into *E.coli (DE3*) (data not shown). The expression and the solubility of the recombinant proteins were tested by 12% SDS-PAGE. However, there was only a low soluble expression of AnHSD in the crude enzyme solution when *E.coli*-AnHSD was expressed alone (Fig.3B Lane 2-3). Next, we co-expressed AnHSD with pGro7 and found soluble protein levels dramatically improved (Fig.3B Lane 4-5). SDS-PAGE analysis also indicated an estimated molecular subunit mass of approximately 36.5 kDa (included an N-terminal 6 × His tag).

**Figure 3.**
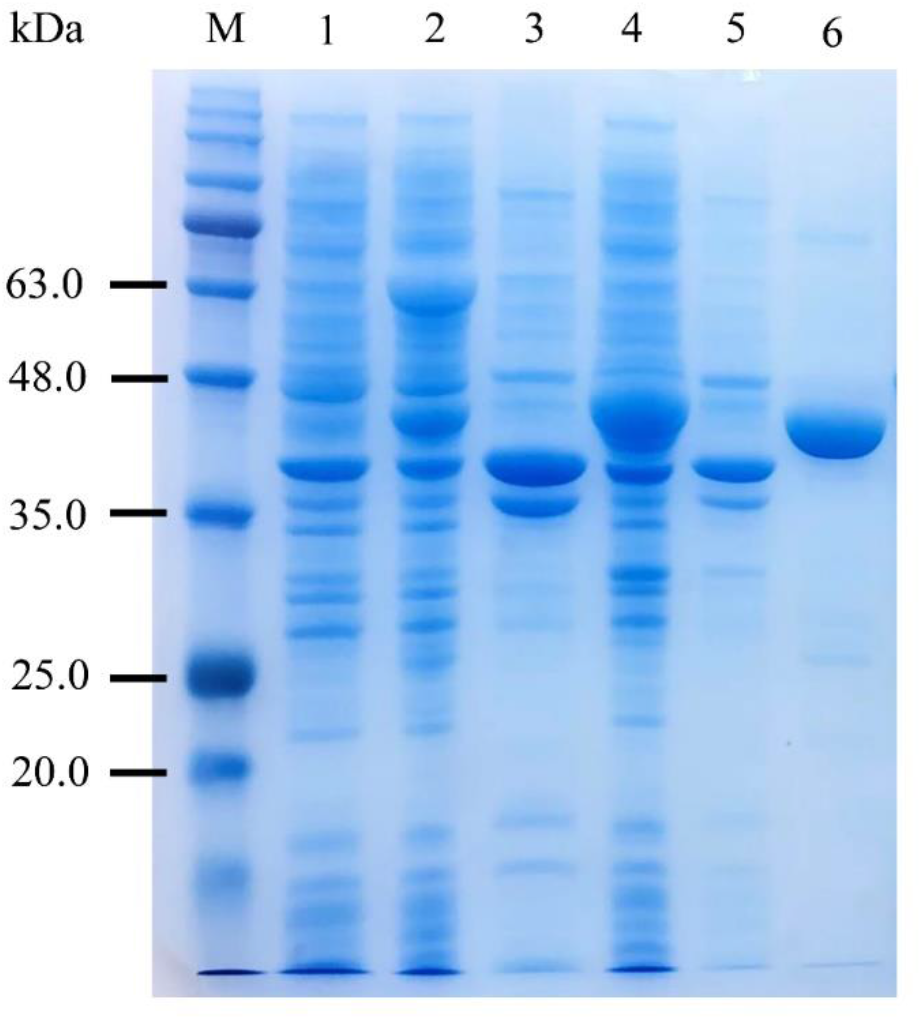
Expression of AnHSD, M: protein marker, Lan1: expression of pRSFDuet-1 vector as control (supernatant), Lane2: the supernatant of pRSFDuet-AnHSD. Lane3: the pellet of expression of pRSFDuet-AnHSD. Lane4: the supernatant expression of pRSFDuet-AnHSD-pGro7. Lane5: the pellet of expression of pRSFDuet-AnHSD-pGro7. Lane6: the purified AnHSD.

Compared with the control group and the experimental group showed a peak of L-ASA (Fig.4 A and B). Mass spectral detection of L-Hse and L-ASA was performed in a targeted MS/MS mode with positive (ESI+) ionization mode. The activity catalytic of AnHSD was further demonstrated from that the formation of L-ASA ([M + H] ^+^ = 118) from L-Hse ([M + H] ^+^ = 120) (Fig.4 C and D).

**Figure 4.**
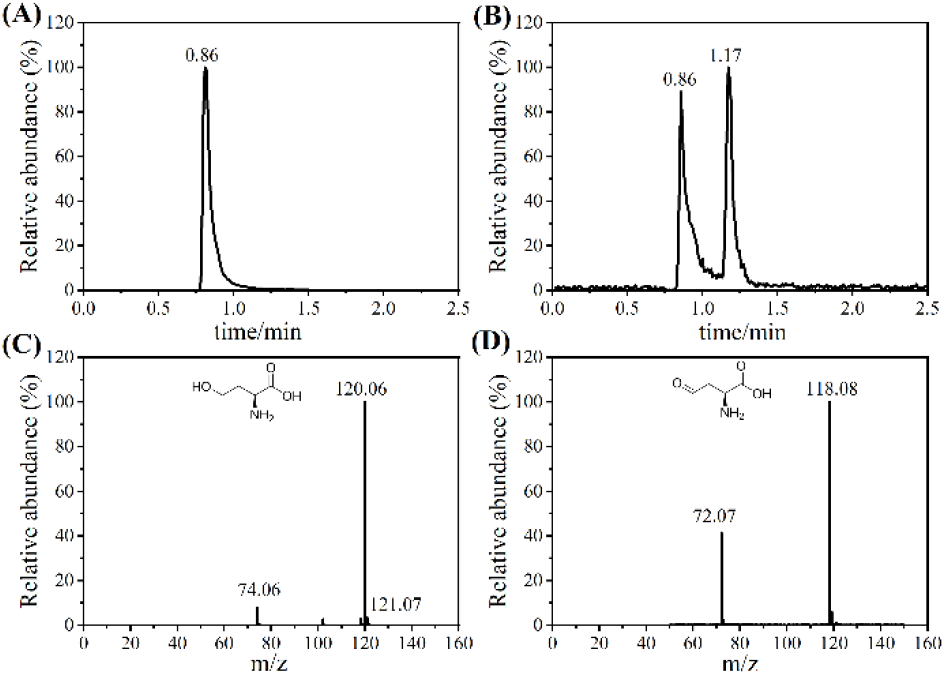
LC-MS/MS chromatograms showing synthesis L-ASA after incubating L-Hse with AnHSD. Peaks at 0.86 and 1.17 min corresponded to the stands L-Hse and L-ASA, respectively. (A) The products from free-enzyme controls only detected L-Hse. (B) The products from AnHSD samples detected not only L-Hse but also L-ASA. (C) ESI-MS detection of L-Hse. (D) ESI-MS detection of L-ASA.

### Characterization of purified AnHSD properties

The effect of temperature on AnHSD activity was shown in Fig.5A, the optimal temperature profile of AnHSD showed that the activity increased with temperature until its peak at 40 °C, but decreased rapidly when the temperature was above 40 °C, and most of the enzyme activity was lost at 70 °C. The pH profile showed a bell-shape (Fig. 5B). It was also shown that the AnHSD activity improved significantly with the pH ranging from 5.0 to 10.0, and achieved optimal activity at pH10.0. The residual enzyme activity was more than 60% at pH 12.0. The stability of temperature and pH on AnHSD were determined at the same time. As shown in Fig. 5C, the temperature stability test for recombinant AnHSD showed that the enzyme was comparative stable at 20 −40 °C, the remaining enzyme activities were no less than 90%. After that, the enzyme activity decreased all the time and lost almost activity at 70 °C. Fig. 5D showed that AnHSD possessed worse stability at acidic pH and had high stability at pH10.0. The enzyme was inactive after 1 h incubation at pH 12.0.

**Figure 5.**
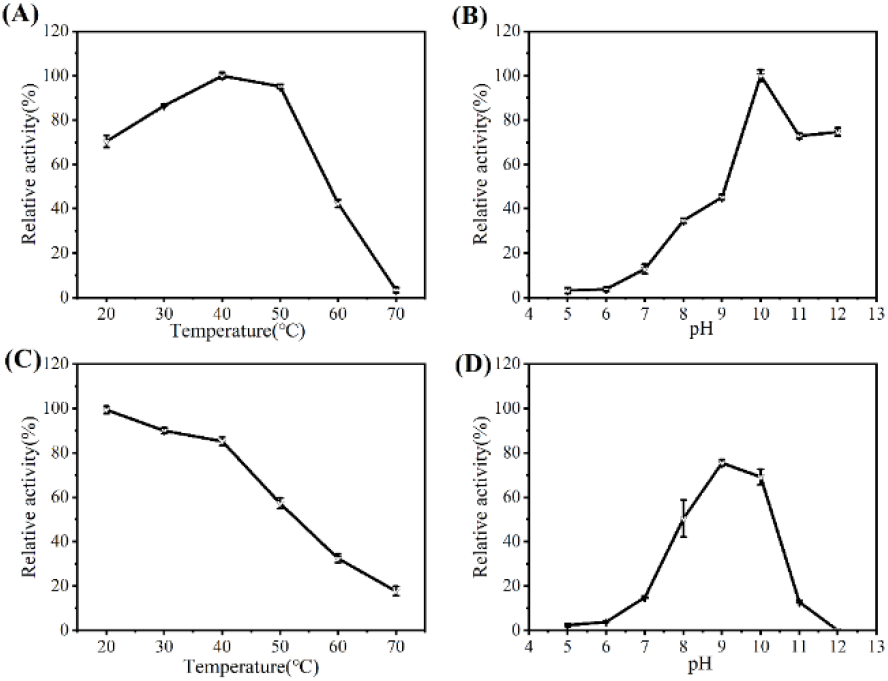
(A) Effects of temperature on AnHSD activity and the maximum activity (41.36 μmol·min^-1^·mg^-1^) was set to 100%. (B) Effects of pH on AnHSD activity and the maximum activity (40.68 μmol·min^-1^·mg^-1^) was set to 100%. (C) Effects of temperature on AnHSD stability. (D) Effects of pH on AnHSD stability.

In a further set of experiments, we investigated the influence of different metal ions on AnHSD activity (Table 1). Most transition metals decreased the specific activity of AnHSD. In particular, Co^2+^ and Zn^2+^ (<1 mM) completely inhibited the activity of AnHSD. DTT and 2-mercaptoethanol had little negative influence on AnHSD, and increasing the levels of disulfide reductants could enhance the activity of AnHSD levels to some extent. As shown in Table 2, we determined the influence of 20 amino acids on AnHSD activities. Notably, L-Cys strongly inhibited AnHSD activity. In addition, the lower concentrations (1 mM) of L-Met, L-Trp stimulated AnHSD activity, and higher concentrations (10 mM) had a certain inhibitory effect. Under the stimulation of L-Lys, L-Pro or L-Arg, a significant increase in AnHSD activity was observed.

**Table 1.** Effect of metal ions on AnHSD activity

| Metal ions | Relative activity (%) |  |
| --- | --- | --- |
|  | 0.1mM | 1mM |
| Control | 100 ± 1.45 | 100 ± 1.23 |
| Mg<sup>2+</sup> | 80.41 ± 0.75 | 72.93 ± 3.33 |
| Zn<sup>2+</sup> | 35.81 ± 2.22 | 8.14 ± 3.9 |
| Mn<sup>2+</sup> | 72.57 ± 1.31 | 5.35 ± 2.49 |
| Co<sup>2+</sup> | 76.11 ± 2.38 | 0.00 ± 1.28 |
| Ni<sup>2+</sup> | 78.77 ± 3.99 | 75.94 ± 1.78 |
| Ca<sup>2+</sup> | 81.59 ± 2.76 | 81.11 ± 1.29 |
| Cu<sup>2+</sup> | 81.77 ± 0.82 | 66.25 ± 8.81 |
| Fe<sup>2+</sup> | 82.11 ± 3.24 | 75.79 ± 2.09 |
| Fe<sup>3+</sup> | 90.51 ± 2.95 | 63.06 ± 5.92 |
| DTT | 91.31 ± 3.99 | 96.42 ± 5.44 |
| EDTA | 82.26 ± 2.63 | 75.52 ± 3.89 |

**Table 2.** Effect of amino acids on AnHSD activity

| Amino acids | Relative activity (%) |  |
| --- | --- | --- |
|  | 5 mM | 10 mM |
| Control | 100.30 ± 1.46 | 100.43 ± 1.27 |
| L-Cys | 19.80 ± 2.60 | 7.71 ± 6.99 |
| L-Gly | 80.50 ± 10.23 | 96.04 ± 5.62 |
| L-His | 81.78 ± 2.42 | 81.47 ± 4.27 |
| L-Val | 84.88 ± 3.49 | 94.27 ± 6.04 |
| L-Ser | 88.30 ± 3.24 | 86.44 ± 2.63 |
| L-Asp | 89.24 ± 2.74 | 76.69 ± 1.53 |
| L-Tyr | 91.17 ± 25.58 | 92.81 ± 7.08 |
| L-Ala | 94.06 ± 6.05 | 105.85 ± 5.74 |
| L-Ile | 96.53 ± 2.87 | 81.44 ± 2.20 |
| L-Leu | 96.74 ± 5.27 | 107.65 ± 5.08 |
| L-Thr | 99.21 ± 4.94 | 105.00 ± 6.55 |
| L-Gln | 99.66 ± 1.90 | 89.37 ± 2.83 |
| L-Asn | 99.91 ± 1.05 | 90.52 ± 4.92 |
| L-Phe | 100.67 ± 1.41 | 100.85 ± 4.84 |
| L-Glu | 101.71 ± 6.99 | 99.42 ± 6.78 |
| L-Met | 105.58 ± 4.77 | 98.81 ± 0.62 |
| L-Trp | 105.91 ± 7.61 | 93.84 ± 1.92 |
| L-Lys | 108.07 ± 9.75 | 117.52 ± 2.66 |
| L-Pro | 110.45 ± 6.15 | 104.78 ± 0.88 |
| L-Arg | 119.38 ± 10.65 | 117.40 ± 2.49 |

### AnHSD substrate specificity analysis and kinetic properties

A variety of alcohols were used as substrates for determination of the catalytic capacity of purified AnHSD. These results directly demonstrate that AnHSD showed weak oxidation activity towards 1-butanol and isopropyl alcohol, the specific activities were 3.92 ± 0.10 μmol·min^-1^·mg^-1^ and 2.64 ± 0.53 μmol·min^-1^·mg^-1^, respectively, but without any notable activities to residual alcohols, including 2-aminoethanol, 4-amino-1-butanol, 2-phenylethanol, 1,2-propanediol, 1,3-butanediol, 2-methyl-1-propanol, 3-methyl −1-butanol, 2-methyl-2-propanol, 2-butanol, 2-methyl-1-butanol, 3-aminopropanol.

The kinetic parameters of AnHSD were measured by plotting the initial velocity against the L-Hse concentration. The values of *V_max_*, *K_m_*, and *K_cat_* were found to be 180.70 ± 10.35 μmol·min^-1^·mg^-1^, 6.30 ± 1.03 mM, and 462.71s^-1^, respectively. Therefore, the catalytic efficiency (*K_cat_*/*K_m_*) was 73.44 s^-1^·mM^-1^. With respect to the cofactor usage, AnHSD showed the highest activities with NAD^+^ as the electron receptor. A reasonable activity was, nevertheless, not obtained on NADP^+^.

## Discussion

As a known oxidoreductase, HSD can catalyze the reversible conversion of L-ASA to L-Hse with a nucleotide cofactor-dependent reduction reaction, generated L-Met, L-Ile, L-Thr (3, 18), constituted a key metabolic hub in the metabolic pathways of L-aspartate family amino acids (AFAAs). However, recent studies indicated that some microorganisms, such as *A. nicotinovorans* and *R. leguminosarum*, could utilize L-Hse as a sole carbon source for normal survival and metabolism to acclimate to a homoserine-rich environment (12, 19). Although several strains capable of degrading L-Hse have been studied, the molecular mechanism and gene responsible for this degradation have not been reported. Our study discovered a putative protein, denoted as AnHSD, was responsible for the metabolism of L-Hse in *A. Nicotinovorans*.

The phylogenetic tree clearly revealed the evolutionary relationship between AnHSD and other related HSDs (Fig.2). Two HSDs from *A. nicotinovorans*, one was AnHSD and another was AnHSD-109 (WP_055972109.1), were assigned to two separate families. AnHSD-109 was homologous to other HSDs from the aspartate metabolic pathway used to synthesize L-Thr, L-Met, and L-Iso (20). In striking contrast, AnHSD had a quite similarity with AnHSD-109 only 33.96%, also was distinct from other mono- and bi-functional HSDs, but clearly divided into an independent new HSDs family, none of these have been characterized concerning biocatalysis. This discrepancy was likely due to the different physiological ways of making use of L-Hse in various creatures (12, 21). This situation was also present in *R. leguminosarum* which was in the same evolutionary branch of *A. nicotinovorans*. Two HSDs from *R. leguminosarum* also were located in distinctly different branches of the evolutionary tree (Fig.2) and they possessed low sequence identity (35.45%). A microarray study of gene expression indicated that one of putative homoserine dehydrogenase gene on *R.leguminosarum* was specifically up-regulated in the pea rhizospheres with rich L-Hse (10, 22). This suggests that RlHSD (WP_011654697.1) might have a role in the degradation of L-Hse. A number of genes around the *rlhsd* gene were homologous to ABC-type importers, suggesting that they were likely required for transport of L-Hse into the cytoplasm (11). The results presented above illustrated that these strains have an alternate genetic pathway for homoserine catabolism. Same as *rlhsd* discussed above, some genes around *anhsd* were homologous to ABC-type importers and the cation transport regulator ChaB that could provide metal ion support for dehydrogenase. The presence of AnHSD suggests that HSD of A. might be involved in the degradation of L-Hse to L-ASA More critically, there was also a putative aldehyde dehydrogenase (AlDDH), which was likely to continue to catalyze the formation of L-Asp from L-ASA specifically, leading to L-ASA entered the aspartate metabolic pathway or the TCA cycle pathway again (Fig.6B).

**Figure 6.**
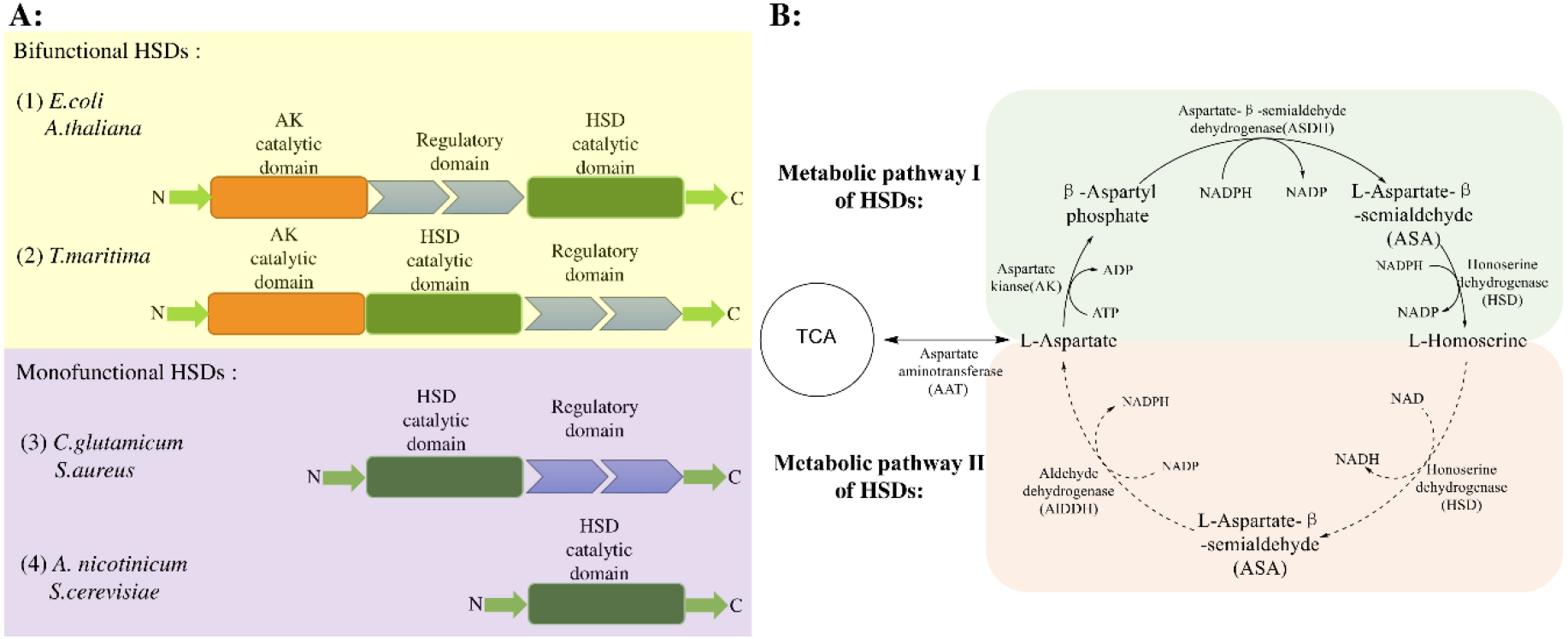
A. Orientation of the different domains in bifunctional AK-HSDs and monofunctional HSDs. B. Two types of metabolic pathways of HSDs. (I) represents the main aspartate metabolic pathway. (II) represents the predicted alternative utilization pathway of L-Hse.

To understand better the function of AnHSD, we undertook a systematic characterize the dynamics of AnHSD. AnHSD catalyzed the oxidation of L-Hse to L-ASA with higher catalytic efficiency, however, no corresponding reduction activity of L-ASA was detected. The *K_cat_/K_m_* of AnHSD oxidation activity could reach 3.86 s^-1^·mM^-1^ under the conditions of 40 °C, pH10.0, and 1mM NAD^+^. The oxidation activity of AnHSD was barely detectable when used NADP^+^ as coenzyme. With respect to the cofactor usage, AnHSD showed the highest activities with NAD^+^ as electron receptor. Numerous studies have reported HSDs were dual coenzyme dependent (3, 23). One must note that PhHSD from *Pyrococcus horikoshii*, had a strong preference for NAD^+^, but NADP^+^ as a strong inhibitor of NAD-dependent oxidation of PhHSD (24, 25). The most striking difference between NAD^+^ and NADP^+^ was the C2 phosphate group of adenine ribose, thus the amino acid residues that interact with the C2 region were considered to be the key residues responsible for the specificity of the coenzyme (26). NAD(P)^+^ binding sites have a β-α-β motif that constituted the conserved sequence of GXGXXG/A (27). The major determinant of NAD^+^ specificity was the presence of an aspartate residue, which formed double-hydrogen bonds to hydroxyl groups located on both the C2 and C3 positions in the ribosyl moiety of NAD^+^ and created a negative electrostatic potential to the binding site. Usually, this residue in NADP-dependent HSDs was replaced by a smaller and uncharged residue such as Gly, Ala, and Ser, and formed a positive binding pocket for the 2’-phosphate group of NADP^+^ with surrounding amino acids, including Asp and Lys residues (28). Indeed, the alignment shown in Fig.1A appears to confirm this theory. All double NAD(P)-dependent HSDs have a G at the indicated position, while the single NAD-dependent HSDs have a N in the novel HSDs family. The crucial Asn in AnHSD could interact with the hydroxyl groups at the C2 and C3 positions of NAD^+^ adenine ribose to produce stable hydrogen bonds (29), as well as occupied a significant portion of the space of the C2-phosphate group of NADP, leading to that AnHSD cannot take advantage of NADP as a coenzyme or made use of NADP inefficiently. It is worth note that Ala was also present at *L.musarum* in the novel HSD family, LmHSD may tend to specifically use NADP as a sole coenzyme carrier. We speculate that the novel HSDs family may have the characteristics of single coenzyme utilization.

HSD as a crucial enzyme in metabolic pathways of AFAAs and branched-chain amino acids (BCAAs), correspondingly was suppressed by L-threonine, L-lysine, and L-methionine (30). In the threonine-sensitive AK-HSD, the linker region between AK and HSD was responsible for the regulatory effect of L-threonine (Fig.6A) (31–33). In the middle region of *Arabidopsis Thaliana* AK-HSD, two potential Threonine binding sites have been identified. The combination of Gln443 and Threonine led to the inhibition of AK activity, this situation contributed to the bind of Threonine to the second Gln524, resulted in the suppression of HSD activity (34). The arrangement of AK and HSD in *Thermus thermophilus* was not the same as AK-HSD from *E. coli* and *A. thaliana*, the domains lined up in the order HSD, AK, and regulatory domain (Fig.6A). Gln524 in *A.thaliana* AK-HSD was replaced by Ala709 and generated threonine-insensitive AK-HSD (35). The monofunctional CgHSD and SaHSD belonged to the ACT domain-containing protein family and were strongly inhibited by its end-products L-threonine and L-isoleucine (14, 33, 36). However, some studies have shed light that monofunctional ScHSD (from *Saccharomyces cerevisiae*) and GmHSD (from soybean) were naturally occurring feedback resistant enzyme for aspartate-derived products (which at physiological levels) due to the absence of ACT-domain (37–39). Thence, the feedback inhibition of amino acids synthesis in *yeast* was mainly concentrated on AK (HOM3) (40). In the next study, we examined the effect of the 20 amino acids on AnHSD. Combining extensive biochemical analyses with the sequence characterization, we identified that AnHSD was a novel monofunctional HSD that had no extension of the C-terminal ACT domain, and the activity of AnHSD was not affected by L-threonine, L-lysine, L-methionine at 10 mM concentration (Table 2). The above results illustrated that AnHSD was a natural feedback resistant enzyme such as ScHSD and GmHSD. Remarkably, L-aspartate showed slight inhibition (reduced to ~70% that of the original activity) to the activity of AnHSD. It was currently hypothesized that too much L-ASA were synthesized by AnHSD and subsequently transformed to L-aspartate by AlDDH. Thus, the feedback resistance was caused by excessive L-aspartate and was responsible for maintaining the balance between L-Hse oxidation pathway and the main metabolic pathway of aspartate amino acids (Fig.6B).

Detailed analysis of L-cysteine inhibition of AnHSD (Table 2) showed L-cysteine to be a competitive inhibitor *versus* L-Hse (*K*_i_=0.089mM). Several studies have reported the role of L-cysteine in affecting the activity of StHSD. The sulfur atom of L-cysteine was covalently combined to the nicotinamide ring could inhibit the activity of StHSD in an enzyme-NAD-cysteine complex manner (41). As in StHSD studies, the inhibition of AnHSD activity induced by L-cysteine was probably because that the irreversible covalent binding between L-cysteine and the key amino acids at the active site.

Some studies have revealed the impact of K^+^ and Na^+^ on HSDs. K^+^ was an activator and Na^+^ was an inhibitor for *E.coli* AK-HSDs (42). However, K^+^ and Na^+^ both were an activating agent for ScHSD (43). In the course of determining AnHSD activity, we found that K^+^ and Na^+^ both displayed activation function for AnHSD and potentiated the oxidation function of AnHSD when increased concentration of K^+^ and Na^+^. We also tested the effect of the buffer system on AnHSD reaction. The results showed that phosphate buffer (including potassium phosphate and sodium phosphate buffer) have a significant effect on AnHSD while the activity of AnHSD was undetected in Tris-HCl buffer which without any metal ions. The results described above demonstrated that positive monovalent cations had an acute influence on AnHSD activity. One possible reason that may explain this finding was that the metal ion was located precisely at the junction between the nucleotide-binding region and the dimerization region, the presence and the size of a metal ion may change the conformation of this part of the protein structure, which in turn affect AnHSD activity (7, 43). In addition, we also found that AnHSD was highly sensitive to Zn^2+^, Mn^2+^, and Co^2+^ and lost the whole activity at a high concentration of metal ions (Table 1), which indicated that the important role of the sulfhydryl group on AnHSD (44). The remaining bivalent metal ions showed slight inhibition on AnHSD, as well as metal chelating agent EDTA had no significant effect on AnHSD (more than 75% at 10 mM). The above studies indicated that the oxidation activity of AnHSD did not rely on bivalent metal ions (45).

Until now, there is no report about genes showing an HSD activity converting L-Hse to L-ASA experimentally in this novel class HSD of PRK 06270 superfamily. In this study, the gene *anhsd* encoded AnHSD from *A.nicotinophilus* was synthesized after codon optimization. The results revealed that the AnHSD protein is responsible for the degradation L-homoserine in strain *A.nicotinophilus*, catalyzes the oxidation of L-homoserine to L-ASA. Our study reports on the metabolic pathway of L-homoserine and reveals that AnHSD protein is present in diverse bacteria. In addition, the functional studies of AnHSD in *A.nicotinophilus* and other microbial strains are necessary for a better understanding of their metabolic features in terms of making use of L-Hse as the sole carbon source.

## Author statement

CRediT (Contributor Roles Taxonomy) author statement as follows: Xiaoxiang Hu: Writing-Original Draft, Conceptualization, Methodology, Software Investigation. Yajun Bai: Data curation, Methodology. Tai-Ping Fan: Validation, Investigation. Tai-Ping Fan: Validation, Investigation. Xiaohui Zheng: Project administration. Yujie Cai: Supervision, Project administration.

## Materials and methods

### Materials, bacterial strains and vectors

L-Hse, Kanamycin, chloramphenicol, arabinose, isopropyl β-D-thiogalactoside (IPTG), β-nicotinamide adenine dinucleotide (NAD^+^) reductive form, β-nicotinamide adenine dinucleotide phosphate (NADP^+^) reductive form and other required chemicals in this study were of analytical grade and purchased from Sigma-Aldrich (St. Louis, MO, USA). L-ASA was synthesized by the method of Suzanne L. Jacques (43). These preparations were not pure enough to be used as standards for quantitative analysis. Plasmids pRSFDuet-1, pGro7 purchased from Takara (Dalian, China). *E.coli BL21 (DE3*) from Novagen (Shanghai, China) was used to express recombinant enzyme. AxyPrep Plasmid Miniprep Kits for plasmid DNA extraction was ordered from Axygen (Suzhou, China).

### Construction of plasmids and strains

For *in-vitro* characterization of the enzymatic properties of AnHSD (accession number is WP_064723327.1) from *A.nicotinophilus*, we synthesized the *anhsd* gene on the basis of codon optimization and cloned into the kanamycin-resistant pRSFDuet-1 vector to constructed pRSFDuet-AnHSD. Then the obtained plasmid was transformed into *E.coli BL21 (DE3*) competent cells. Transformants were selected by kanamycin resistance and validated by diagnosis and sanger sequences. The resulting strain was named as *E.coli*-AnHSD. Co-expression of the pRSFDuet-AnHSD and the chloramphenicol-resistant pGro7 plasmid which expressed molecular chaperone were transformed into *E.coli BL21 (DE3*) cells using a two-plasmid strategy. Transformants were selected under the control of kanamycin and chloramphenicol and validated according to the above methods. The resulting strain was named as *E.coli*-AnHSD-pGro7. The constructed recombinant strains were cultured in LB medium containing 10 g·L^-1^ tryptone, 10 g·L^-1^ NaCl, 5 g·L^-1^ yeast extract. Kanamycin (50 μg·mL^-1^), chloramphenicol (50 μg·mL^-1^), IPTG (40 mmol·L^-1^) and arabinose (10 mmol·L^-1^) were added to the medium when needed.

### Expression and purification of the AnHSDs and protein assays

Single colonies of *E.coli*-AnHSD were picked from agar plate and incubated into test tubes containing 3 mL LB medium with 50 μg·mL^-1^ kanamycin for overnight growth at 37 °C and 200 rpm·min^-1^. Then 1% inoculum of the precultures were inoculated into shake flasks containing 50 mL LB medium with 50 μg·mL^-1^ kanamycin. Cells were grown at 37 °C until optical density (OD_600_) of 0.6 was reached and then induced with 40 mmol·L^-1^ IPTG. Protein was expressed for 20 hours at 20 °C. The protein expression method of double-plasmid *E.coli*-AnHSD-pGro7 recombinant bacteria was the same as above. Furthermore, 50 μg·mL^-1^ chloramphenicol should be added in the process of cell culture, as well as 10 mmol·L^-1^ arabinose should be added in the process of inducing. Cells were collected by centrifugation at 10000 × g for 5 min and the precipitate was resuspension in 20 mM pH 7.4 phosphate buffer saline (PBS). The suspension was broken by sonication on ice and then centrifuged at 10000 × g for 20 min. The supernatant was incubated with BeaverBeads^™^IDA-Nicker (BEAVER biomedical, China) for 30 min at 4 °C, and the tag-free proteins were then eluted by 50 mM imidazole, the protein was eluted by 250 mM imidazole and then immediately desalted into the desalting buffer. The quality of purified proteins was analyzed by 12% sodium dodecyl sulfate-polyacrylamide gel electrophoresis (12% SDS-PAGE). The total protein concentration was determined using the Bradford method (Solarbio, China) (46).

### Enzyme activity assays

The enzyme activity of AnHSD was determined by monitoring the absorbance of NAD(P)H at 340 nm by UV-6000 METASH spectrophotometer instrument according to the method described previously (24). The standard reaction system of AnHSD for L-Hse oxidation (3.0 mL, 40 °C) containing 20 mM PBS (pH 9.2), 50 mM L-Hse, 150 mM KCl, 1 mM NAD^+^ and 50 μL purified AnHSD. The assay mixture for L-ASA reduction contained the following components in a total volume of 3.0 mL: 20 mM PBS (pH 6.0), 10 mM L-ASA, 0.12 mM NAD(P)H, 50 μL purified AnHSD. The reaction solution without enzyme was used as a control to explain any spontaneous hydrolysis of L-Hse or L-ASA. One unit of activity was defined as that 1 μmol NAD(P)H consumed per minute, and specific activity was reported as units per mg protein (extinction coefficient was 6.22 mM^-1^ cm^-1^).

The reaction products of AnHSD were extracted and detected by LC-MS/MS operated in multiple reaction monitoring (MRM) mode with WATERS ACQUITY UPLC (Waters, UK). A BEH C18 analytical column was used to separate various components of samples. The mobile phase consisted of solvent A (0.1% formic acid) and solvent B (100% acetonitrile), the condition of the mobile phase was as follows: Solvent A (in%): 100 (40min), 70 (45min), 20 (50min), 100 (55min); Solvent B (in%): 0 (40min), 30 (45min), 80 (50min), 0 (55min). The flow rate, column temperature, and injection volume were set to 0.3 mL·min^-1^, 45 °C, and 1 μL.

### Characterization of enzymatic properties of purified AnHSD

Substrate specificity for a large number of alcohol compounds was tested with the purified AnHSD according to the enzyme activity determination described above. The substrates were included 2-aminoethanol, 4-amino-1-butanol, 2-phenylethanol, 1,2-propanediol, 1,3-butanediol, 1-butanol, 2-methyl-1-propanol, 3-methyl −1-butanol, 2-methyl-2-propanol, 2-butanol, isopropanol, 2-methyl-1-butanol, 3-aminopropanol.

The effects of temperature on AnHSD were measured over a range of 20-70 °C in 20 mM PBS, pH7.2. To investigate the effect of pH on AnHSD, the enzyme assay was carried out at the pH ranging from 3.0 to 12.0 at 40 °C. The maximum activity measured in the determination process was set to 100%. To study the influence of temperature and pH on the stability on AnHSD, the enzyme was determined using standard assay reaction incubated 1 hour at a temperate range from 20 to 70 °C and a pH range of 3-13. The enzyme activity before pre-incubation was set to 100%.

To evaluate the influence of metal ions on AnHSD activity, 0.1 mM and 1 mM (final concentration) of Ca^2+^, Mg^2+^, Zn^2+^, Cu^2+^, Fe^2+^, Fe^3+^, Mn^2+^, Co^2+^, Ni^2+^, metal chelating agent EDTA or reductants DTT were added to the enzyme solution, and the residual activity was measured.

The effects of 20 amino acids were verified by adding 5 mM and 10 mM (final concentration) different amino acids solutions (L-Gly, L-Ala, L-Val, L-Leu, L-Ile, L-Met, L-Trp, L-Phe, L-Pro, L-Ser, L-Thr, L-Cys, L-Tyr, L-Asn, L-Glu, L-Asp, L-Gln, L-Lys, L-His, L-Arg) to the reaction mixtures.

### Determination of kinetic parameters of the AnHSD with single substrate

According to the Lineweaver-Burk diagram, steady-state kinetic (*K_m_*, *V_max_* and *K_cat_*) of the purified AnHSD were estimated with L-Hse at concentrations ranging from 0 to 30 mM under optimal temperature and pH conditions. The kinetic parameters were evaluated by nonlinear fitting using Origin 2019 software.

### Sequence analysis

The BLAST program (https://blast.ncbi.nlm.nih.gov/Blast.cgi) was used for AnHSD gene homology searches, for predicting homologs with the most closely related sequences from other sources in the NCBI database. The phylogenetic trees were constructed using the MEGA7.0. Protein sequence alignments were obtained with the online server ClustalW (https://www.genome.ip/tools-bin/clustalw) and ESpript3.0 software. The InterPro tool was used for predicting the presence of domains and classing it into families (47).

## Declaration of competing interest

The authors declare that they have no competing interests.

## Author contributions

The manuscript was written through contributions of all authors. All authors have given approval to the final version of the manuscript.

## Acknowledgements

We thank the National Key Scientific Instrument and Equipment Development Project of China (2013YQ17052504), the Program for Changjiang Scholars and Innovative Research Team in the University of Ministry of Education of China (IRT_15R55), the seventh group of Hundred-Talent Program of Shanxi Province (2015), and The Key Project of Research and Development Plan of Shaanxi (2017ZDCXL-SF-01-02-01) for financial support.

